# Withdrawal-induced escalated oxycodone self-administration is mediated by kappa opioid receptor function

**DOI:** 10.1101/177899

**Authors:** Jacques D. Nguyen, Dean Kirson, Michael Q. Steinman, Reesha Patel, Sophia Khom, Florence P. Varodayan, David M. Hedges, Christopher S. Oleata, Yanabel Grant, Marisa Roberto, Michael A. Taffe

## Abstract

**Background:** Prescription opioid addiction is a significant health problem characterized by compulsive drug seeking, withdrawal and chronic relapse. We investigated the neurobiological consequences of escalation of prescription opioid use using extended access to intravenous oxycodone self-administration in rats.

**Methods:** Male Wistar rats acquired oxycodone self-administration (0.15 mg/kg/infusion, i.v.) in 1h or 12h access sessions. Electrophysiological and immunohistochemical studies investigated the effects of oxycodone self-administration on kappa opioid receptor (KOR) regulation of GABAergic signaling and dynorphin expression in the central nucleus of the amygdala (CeA).

**Results:** Rats given 12h access to oxycodone for 5 sessions/week (LgA) escalated their responding more than rats given 1h oxycodone (ShA) or 12h saline access. Slowed escalation of responding was found in rats given 12h access for 7 sessions/week (LgA-7day) or rats pretreated with the KOR antagonist nor-binaltorphamine dihydrochloride (norBNI) before LgA (norBNI+LgA). The KOR agonist U-50488 decreased GABA release in CeA neurons of all groups except LgA. norBNI increased GABA release in control group neurons, suggesting tonic KOR activity. This activity was abolished in ShA, norBNI+LgA, and LgA-7day rat neurons, consistent with decreased CeA dynorphin immunoreactivity observed in LgA-7day rats. However, norBNI effects were reversed (decreased CeA GABA release) in LgA rat neurons.

**Conclusions:** The experience of intermittent extended withdrawal periods accelerates the escalation of oxycodone self-administration and causes greater dysregulation of CeA KOR-mediated GABAergic signaling. A KOR agonist/antagonist switch effect seen with other drugs of abuse was absent, which suggests that oxycodone-induced neuroadaptations may be distinct from those resulting from other drugs of abuse.

## INTRODUCTION

Non-medical opioid abuse is a significant global problem, with an estimated 33 million users of opiates and prescription opioids worldwide (1). Approximately 2 million people in the US have a prescription opioid related abuse disorder (2), which may increase the likelihood of later illicit opioid use (3), and prescription opioid related overdose deaths have drastically increased over the last two decades (4). Despite the growing impact of prescription opioids on public health, few pre-clinical studies have investigated self-administration of oxycodone, one of the most commonly prescribed medications (OxyContin^®^ or as part of Percocet^®^). These studies have shown that oxycodone self-administration causes behavioral changes (5), and sometimes physical dependence and withdrawal (6) in mice. Oxycodone self-administration is similar between male and female rats during early stages of training (7), and male rats trained to self-administer oxycodone under extended access conditions (12h) exhibited a progressive escalation of drug intake (8) similar to heroin escalation under similar 12h access conditions (9, 10). Mice also exhibit escalation of oxycodone self-administration under 4h extended access conditions (11).

The negative reinforcement hypothesis advanced to explain escalating drug intake under extended-access conditions (12, 13) holds that a dysphoric or negative affective state that is experienced during the daily withdrawal from drug intravenous self-administration (IVSA) grows increasingly severe with sequential sessions. Indeed, brain reward thresholds progressively increase with self-administration of heroin in 23h but not 1h daily sessions (14) and correspondingly the somatic signs of withdrawal from heroin increase progressively from 12 to 48h in adult rats (15). The negative affective state associated with chronic exposure to drugs of abuse is thought to be mediated in part by dynorphin/kappa opioid receptor (KOR) signaling (16-18). Increased KOR activity is aversive and KOR antagonists are currently being investigated as potential treatments for stress-induced drug-seeking behaviors and withdrawal-related anhedonia (19). Intra-cerebroventricular administration of the long lasting KOR antagonist, nor-binaltorphimine (norBNI), attenuates stress-induced changes in reward threshold following 20h withdrawal from cocaine (20). Systemic or local administration of norBNI in the basolateral amygdala also reverses stress-induced neuroadaptations mediating reinstatement of cocaine-seeking (21) or nicotine preference (22). Microinfusions of norBNI in the central nucleus of the amygdala (CeA) block repeated cocaine exposure-induced locomotor sensitization and attenuate the heightened anxiety-like behavior observed during withdrawal from cocaine self-administration in the defensive burying paradigm (23).

The CeA is involved in the processing of internal and external emotional stimuli related to anxiety and stress (24) and thus is a critical mediator of escalated drug intake. The CeA is a primarily GABAergic nucleus that also contains numerous stress-related neuropeptides and serves as the major output region of the larger amygdaloid complex. CeA synapses are highly sensitive to many drugs of abuse (12, 25) as functional neuroadaptations in the CeA have been observed following exposure to alcohol (26), cocaine (27), methamphetamine (28), nicotine (29), and opioids (30, 31). The normal dynorphin/KOR inhibition of GABAergic transmission becomes dysregulated in the CeA following escalated cocaine IVSA (23), suggesting that alteration of CeA dynorphin/KOR-mediated signaling may be directly involved in the escalation of drug self-administration under extended access conditions. Relatedly, long-lasting blockade of KOR via systemic administration of norBNI attenuates the escalation of IVSA of methamphetamine or heroin (9, 32). These latter findings show that the 12h interval between self-administration sessions in a 12h long-access heroin IVSA paradigm, or 18h in a 6h methamphetamine IVSA paradigm, is sufficient to induce KOR-mediated negative affect due to drug withdrawal. The negative-reinforcement hypothesis suggests that longer-durations of withdrawal from opioid intoxication might be associated with further escalation of drug intake, since the somatic withdrawal peaks after 48h of drug discontinuation (15). Surprisingly this hypothesis has not been tested directly for long-access drug self-administration.

In this study, we characterized the early stages of prescription opioid addiction and determined if the escalation of opioid intake with extended (12h) access to intravenous oxycodone self-administration was affected by abstinence and by re-engagement of drug seeking (re-escalation) following detoxification. We hypothesized that escalation of oxycodone self-administration is mediated by the negative motivational state associated with oxycodone withdrawal, and that dysregulation of KOR modulation of GABAergic signaling in the CeA underlies this negative affective state.

## METHODS AND MATERIALS

### Animals

Male Wistar rats (N=62; Charles River, Raleigh, NC) were housed in a humidity and temperature-controlled (23±1°C) vivarium on 12:12h light:dark cycles. Animals entered the laboratory at 11-14 weeks of age and weighed an average of 410.0±5.56 g at the start of the self-administration study. Animals had *ad libitum* access to food and water in their home cages and were housed in pairs throughout the study. All procedures were conducted in the animals’ scotophase, under protocols approved by the Institutional Care and Use Committees of the Scripps Research Institute and consistent with the National Institutes of Health Guide for the Care and Use of Laboratory Animals (33).

### Intravenous Catheterization

Rats were anesthetized with an isoflurane/oxygen vapor mixture (isoflurane 5% induction, 1-3% maintenance) and prepared with chronic indwelling intravenous catheters as described previously (34-36). Catheter patency was assessed once a week after the last session of the week, and if catheter patency failure was detected, data that were collected after the previous passing of this test were excluded from analysis. See Supplemental Information for additional detail.

### Drugs

Oxycodone HCl was obtained from Sigma-Aldrich (St. Louis, MO). Nor-binaltorphamine dihydrochloride (norBNI) was obtained from the National institute on Drug Abuse (NIDA) Drug Supply. (-)-U-50488 hydrochloride was obtained from Fisher Scientific (Hanover Park, IL). CGP 55845A, DL-2-amino-5-phosphonovalerate (DL-AP5) and 6,7-dinitroquinoxaline-2,3-dione (DNQX) were obtained from Tocris (Ellisville, MO). Tetrodotoxin (TTX) was obtained from Biotium (Hayward, CA). For behavioral experiments, all doses are expressed as the salt and were dissolved in physiological saline (0.9%), and for electrophysiology experiments, all drugs were constituted in artificial cerebral spinal fluid.

### Self-Administration Procedure

Intravenous self-administration was trained using operant techniques as previously described for oxycodone (8) and other drugs (see Supplemental Information for details). The training dose of oxycodone used was 0.15 mg/kg/infusion (∼0.1 ml/infusion). The session duration for the Short Access (ShA) group was 1 h and the Long Access (LgA) training sessions were 12h in duration. Rats received self-administration sessions during weekdays (ShA, LgA, norBNI+LgA) or for 7 days per week (LgA-7day). Following acquisition rats were subjected to randomized dose-substitution conditions under a Progressive Ratio (PR) schedule of reinforcement wherein different per-infusion doses of oxycodone (0, 0.06, 0.15, 0.3 mg/kg/inf) were presented in a balanced order on sequential sessions lasting up to 3h. Additional details on the group training conditions and the PR procedure are provided in the Supplemental Information.

### Electrophysiology

Recordings were performed in neurons from the medial subdivision of the CeA of rats 24h after the final session of self-administration; see Supplemental Information for slice preparation. GABAergic activity was pharmacologically isolated with 20 μM DNQX (to block AMPAR), 30 μM DL-AP5 (to block NMDAR), and 1 μM CGP 55845A (to block GABA_B_R) (23). All drugs were applied by bath superfusion and were only applied once per slice. The methods for intracellular recordings of evoked inhibitory post-synaptic potentials (eIPSPs) and wh*o*le-cell patch clamp recording of miniature inhibitory post-synaptic currents (mIPSCs) are described in the Supplemental Information.

### Immunohistochemistry

Brain tissue was collected from LgA-7day rats (N=7) and from age-matched naïve controls (N=8) and cryosectioned for immunostaining. Sections containing CeA were collected 24h after the final session of self-administration and analyzed for dynorphin A immunoreactivity. See Supplemental Information for additional detail.

### Data Analysis

Analysis of the IVSA data was conducted with repeated-measures Analysis of Variance (rmANOVA) on the number of infusions earned during the acquisition interval (ShA: 1h session; LgA: 12h session) and on the breakpoints reached in the PR study. Within-subjects factors of Session and Drug Dose (PR) and between-subjects factors for Access Duration / Condition were included. Significant main effects were followed with post-hoc analysis using Tukey (within-subjects factors) or Sidak (between-subjects factors) tests for multiple comparisons. Locally evoked IPSP amplitudes were analyzed with Clampfit 10 (Molecular Devices, Sunnyvale, CA). Frequency, amplitude and kinetics of mIPSCs were analyzed and visually confirmed using a semi-automated threshold-based mini detection software (Mini Analysis, Synaptosoft Inc., Fort Lee, NJ). Averages of mIPSC characteristics were based on a minimum time interval of 3 min and a minimum of 50 events. Electrophysiology results did not differ between LgA (5 sessions per week) treatment groups run in different cohorts, thus these data were combined for analysis. Electrophysiological data were analyzed with t-tests or ANOVAs with Bonferroni post-hoc analyses as appropriate, and are presented as mean +/- SEM. All statistics were performed in Prism (Graphpad, La Jolla, CA). In all cases, p<0.05 was the criterion for statistical significance.

## RESULTS

### Escalation of oxycodone self-administration under extended access conditions

The mean number of intravenous oxycodone infusions obtained by rats trained under LgA (N=8) conditions was significantly higher compared to infusions by ShA (N=12) rats (**Fig. 1A**). The rmANOVA confirmed significant main effects of Session [F(14,252)=24.3; *p*<0.0001], of Access Duration [F(1,18)=97.07; *p*<0.0001], and of the interaction of factors [F(14,252)=21.16; *p*<0.0001]. Post-hoc analysis confirmed that significantly more infusions were obtained by LgA rats during sessions 5 and 7-15 compared to session 1, whereas drug intake did not change for the ShA group. LgA rats received significantly more infusions compared to ShA rats during sessions 5-15. Interestingly, analysis of the sessions before and after the 60 h weekend abstinence periods showed that LgA rats significantly increased drug intake following this extended drug deprivation, whereas ShA showed no significant change (**Fig. 1B**). The LgA and ShA rats also exhibited group differences during PR dose substitution (**Fig. 1C**). The rmANOVA confirmed significant main effects of Access [F(1,72)=10.77; *p*<0.01] and of Dose [F(3,72)=3.570; *p*<0.05]. Post-hoc analysis confirmed that significantly more infusions were obtained by LgA rats at oxycodone 0.060 mg/kg/inf compared to ShA rats.

**Figure 1.**
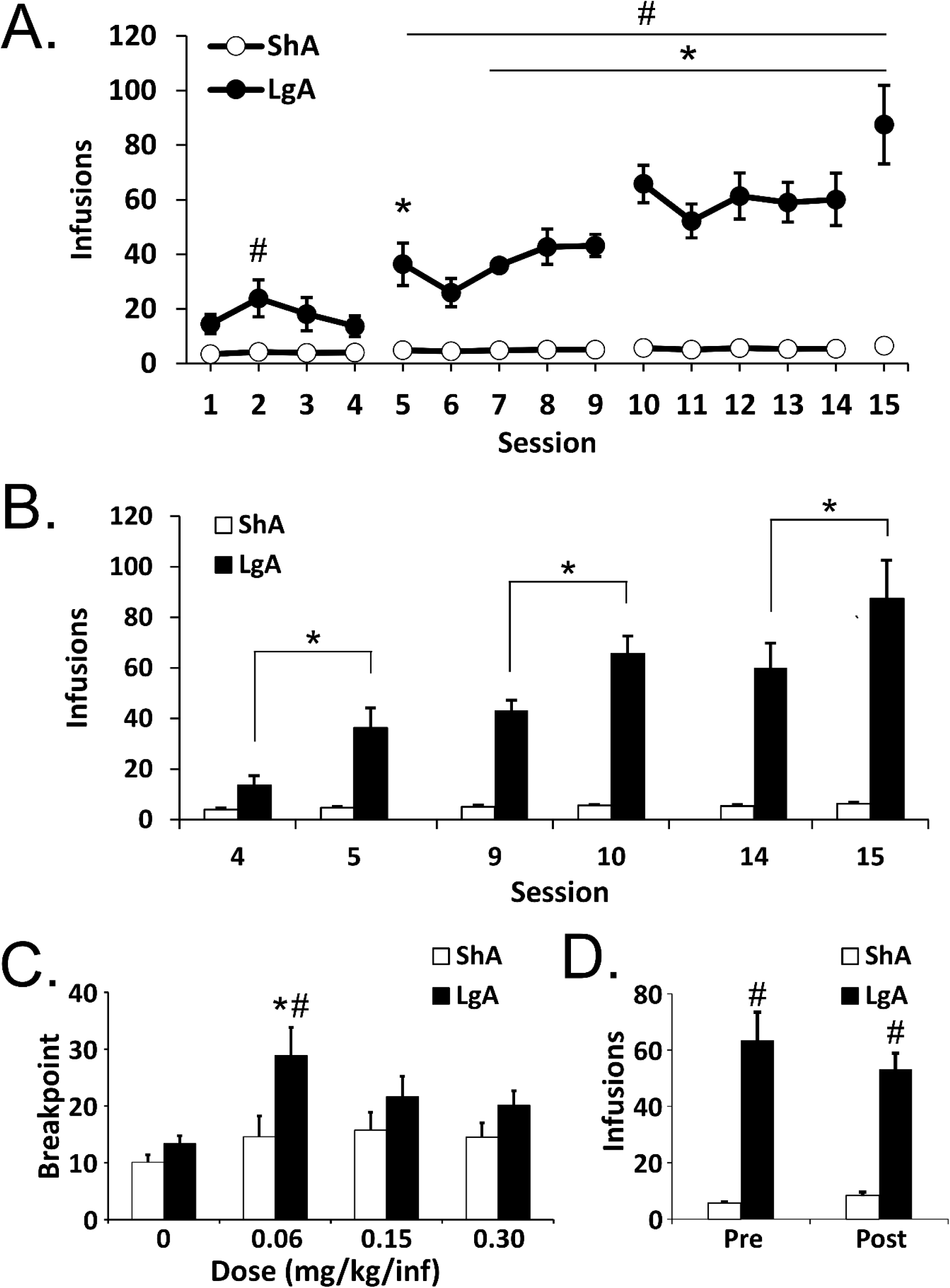
Escalation of Oxycodone Self-Administration. Mean (±SEM) infusions for groups of male rats trained to self-administer of oxycodone (0.15 mg/kg/inf) within Long Access (LgA; 12 h) or Short Access (ShA; 1 h) during A) acquisition, B) pre- and post-weekends, and C) following 30-day abstinence (per 4 sessions). C) Mean (±SEM) breakpoints reached by oxycodone-trained rats under progressive ratio procedure following training. Broken lines between sessions 4 & 5, 9 &10 and 14 & 15 represent 60 h withdrawal period. Significant differences within group from session 1 are indicated by * (unless otherwise indicated) and between group by #.

Following the 30-day abstinence period, neither LgA (N=7) nor ShA (N=12) rats exhibited any significant change in oxycodone infusions (0.15 mg/kg/inf) obtained under a FR1 response contingency (**Fig. 1D**). One rat (LgA) that completed the acquisition interval was not included post-abstinence due to illness. The ANOVA confirmed a significant main effect of Access [F(1,17)=96.26; *p*<0.0001] and the post-hoc analysis confirmed that significantly more infusions were obtained by rats trained under long access compared to short access conditions.

### Effect of short- and long-access oxycodone self-administration on KOR-mediated modulation of GABAergic transmission in the CeA

Electrophysiological studies first examined potential differences in baseline evoked GABAergic transmission in animals with differential experience with oxycodone self-administration. We recorded intracellularly with sharp pipettes from a total of 32 medial CeA neurons (with a mean RMP of -82.06±1.03 mV and a mean input resistance of 175.9±10.2 MΩ) and did not observe any significant differences in basal membrane properties between groups (data not shown). We evoked inhibitory postsynaptic potentials (eIPSP) locally within the CeA. Baseline eIPSP input–output curves generated by equivalent normalized stimulus intensities were similar in CeA from ShA, LgA, and saline control animals (**Fig. 2A**), suggesting no significant changes in baseline evoked GABAergic transmission after oxycodone self-administration.

**Figure 2.**
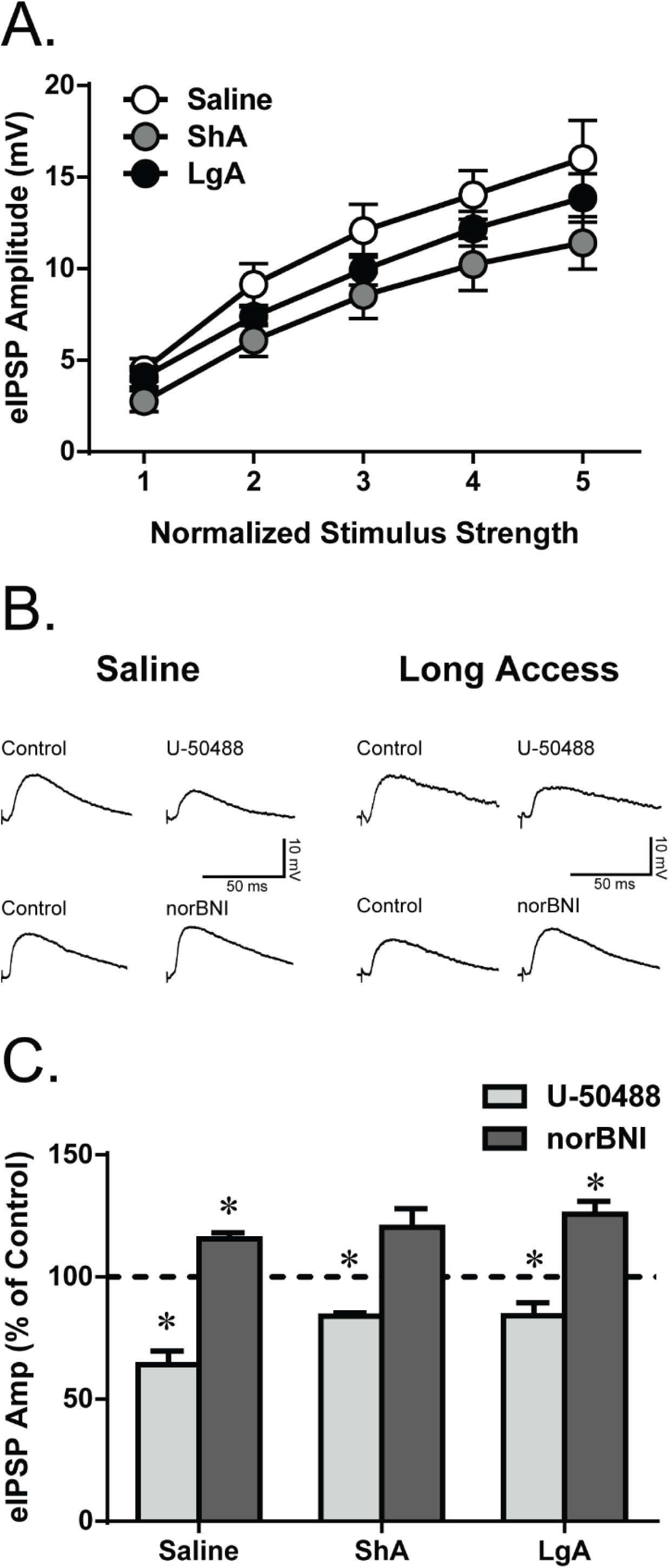
KOR-mediated effects on evoked CeA GABAergic signaling. A) Input/Output baseline evoked inhibitory postsynaptic potential (eIPSP) amplitude relationship for saline, ShA, and LgA animals. B) Representative tracings of eIPSPs recorded before (baseline) and during U-50488 (1 μM) or norBNI (200 nM) superfusion in CeA neurons from saline (Left) and LgA (Right) rats. C) Mean effect of U-50488 and norBNI on eIPSP amplitudes in CeA neurons from saline, ShA, and LgA rats. * denotes significant effect of drug compared to baseline.

We next determined if extended oxycodone access induces any changes in kappa-opioid receptor (KOR)-mediated modulation of GABAergic transmission. We acutely (15 min) applied the KOR selective agonist U-50488 (1 μM) and the KOR antagonist norBNI (200nM) on CeA slices to investigate effects on evoked GABAergic transmission. Notably, U-50488 significantly decreased eIPSP amplitudes (**Fig. 2B & C**) compared to baseline amplitudes [saline: t(3)=6.45, *p*<0.01; ShA: t(3)=12.07, *p*<0.01; LgA: t(6)=2.97, *p*<0.05 for the middle stimulus intensity] in all groups. A one-way ANOVA between groups confirmed significant differences [F(2,12)=4.38; *p*<0.05] in the effects of U-50488, however a Bonferroni post-hoc test failed to confirm any significant differences between individual groups. There was only a trend for U-50488 to have a less pronounced effect in LgA compared to the saline control group [t(12)=2.77, *p*=0.051], indicating that extended access to oxycodone administration may slightly decrease KOR agonist sensitivity. In saline [t(3)=6.09, *p*<0.01 for the middle stimulus intensity] and LgA [t(6)=4.91, *p*<0.01 for the middle stimulus intensity] animals, norBNI caused a significant increase in eIPSP amplitude compared to baseline and a trend [t(3)=2.64, *p*=0.078] towards an increase in eIPSP amplitude in ShA animals. A one-way ANOVA between groups found no significant differences in the effects of norBNI among the saline, ShA, or LgA groups.

We also recorded spontaneous GABAergic miniature inhibitory postsynaptic currents (mIPSC) using whole-cell patch-clamp in the presence of 1 μM TTX to eliminate action potential-dependent release of GABA in the medial CeA of saline control, ShA and LgA (both cohorts) rats. This vesicular form of GABA release is distinct from evoked GABAergic transmission, and can reveal differences in spontaneously active synapses as opposed to effects produced by stimulation of the entire synaptic network (37). We measured the frequency (0.49±0.09, 0.41±0.05, and 0.59±0.09 Hz), amplitude (60.44±6.16, 50.62±5.19, and 68.59±5.68 pA), and kinetics (rise time: 2.29±0.08, 2.57±0.18, and 2.29±0.06 ms; decay time: 5.68±0.59, 7.50±1.09, and 5.57±0.38 ms) of mIPSCs under baseline conditions (**Fig. 3A**) for saline, ShA, and LgA animals respectively. A one-way ANOVA found no significant differences between saline, ShA, or LgA rats for any of the mIPSC parameters, suggesting no significant changes in baseline spontaneous GABAergic transmission after oxycodone self-administration.

**Figure 3.**
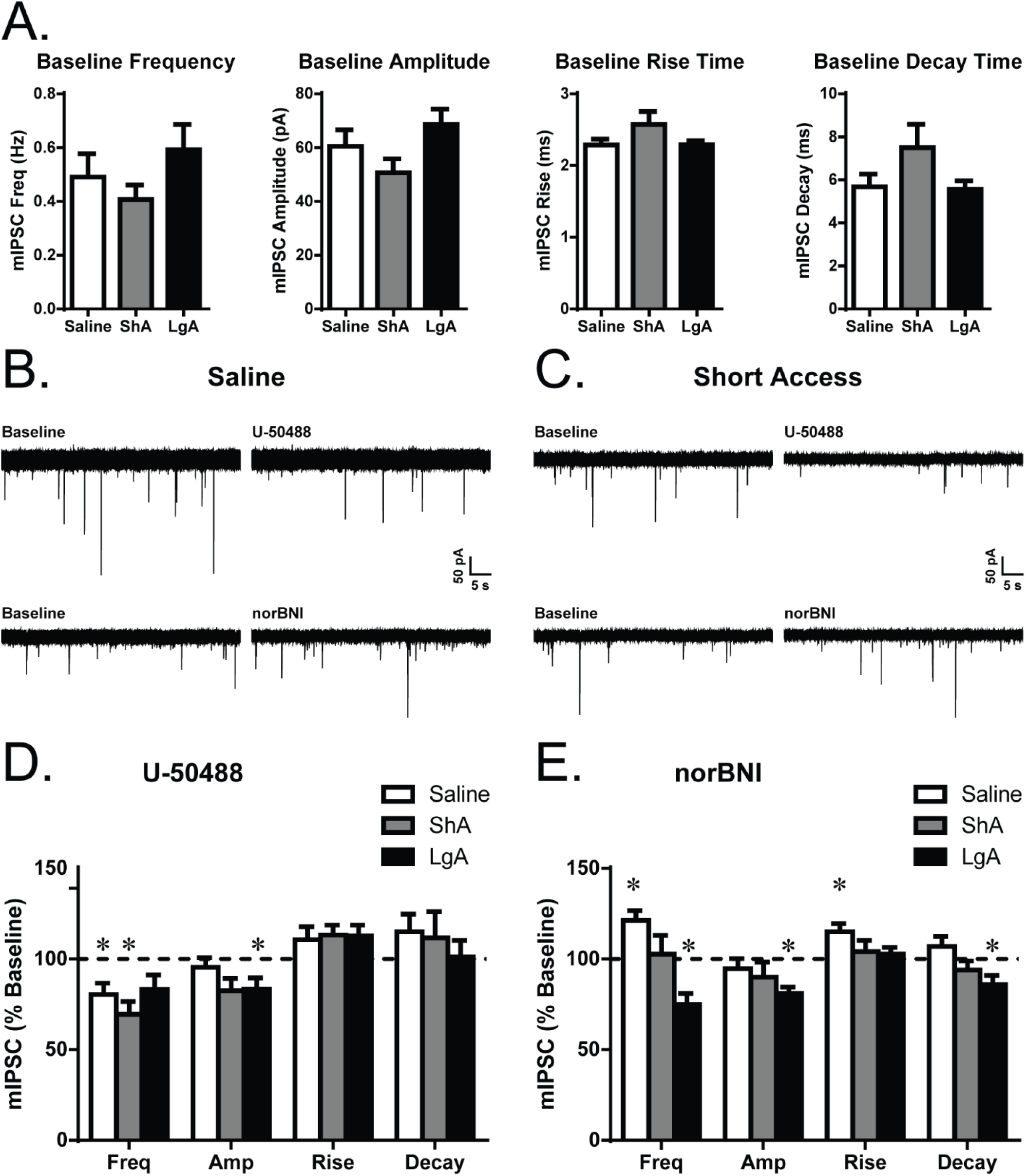
KOR-mediated modulation of CeA spontaneous action potential-independent GABAergic transmission. A) Baseline miniature inhibitory postsynaptic current (mIPSC) frequency, amplitude, and kinetic measurements (rise and decay time) for neurons from saline, ShA, and LgA animals. B) Representative mIPSCs recorded before and after U-50488 (1 μM) or norBNI (200nM) application in neurons from saline and C) ShA rats. Mean effect of D) U-50488 or E) norBNI on mIPSC frequency, amplitude, rise time, and decay time in CeA neurons from Saline, ShA, and LgA rats. * denotes significant effect of drug compared to baseline.

We next assessed the impact of U-50488 and norBNI on mIPSCs (**Fig. 3B-E**). In CeA neurons from saline [t(9)=3.16, *p*<0.05] and ShA [t(5)=4.38, *p*<0.01] rats, U-50488 significantly decreased mIPSC frequency to 80.5±6.17% and 69.59±6.94% of baseline respectively (with no significant changes in amplitude, rise time, or decay time) (**Fig. 3D**), suggesting a decrease in vesicular GABA release. In contrast, in CeA neurons from LgA animals, U-50488 significantly decreased mIPSC amplitudes [t(14)=2.70, *p*<0.05] to 83.46±6.13% of baseline (but not frequency), suggesting a decrease in GABA_A_ receptor function. In CeA neurons from saline control animals, norBNI significantly increased mIPSC frequency [t(7)=3.91, *p*<0.01] and rise time [t(7)=3.30, *p*<0.05] to 121.3±5.45% and 115.0±4.55% of baseline respectively (**Fig. 3E**), supporting a basal KOR receptor-mediated inhibition of GABA release, as observed previously (23). In neurons from ShA rats, norBNI did not alter any of the mIPSC characteristics. In contrast, in neurons from LgA animals, norBNI significantly decreased mIPSC frequency [t(11)=4.17, *p*<0.01], amplitude [t(11)=5.16, *p*<0.001], and decay time [t(11)=2.77, *p*<0.05] to 74.88±6.02%, 80.91±3.70%, and 85.92±5.08% from baseline respectively.

### Blockade of KOR signaling, or uninterrupted daily access, attenuates the escalation of long-access oxycodone self-administration

To further investigate the impact of KOR signaling during acute withdrawal on oxycodone self-administration, a separate cohort of rats was trained under continuous daily sessions (LgA-7day) or administered a systemic intraperitoneal injection of 30 mg/kg norBNI (norBNI+LgA) prior to training 5 days per week. Compared to a second cohort of LgA rats (N=6) trained to self-administer 5 days per week, LgA-7day (N=11) and norBNI+LgA (N=6) rats self-administered significantly fewer drug infusions, e.g., 61.2 (±6.3) and 59.3 (±9.9) during session 17 corresponding to 35.2% and 34.6% less intake, respectively (**Fig. 4A**). The rmANOVA confirmed significant main effects of Drug Condition [F(3,25)=9.49; *p*<0.001], of Time [F(16,400)=18.11; *p*<0.0001], and of the Drug Condition x Time interaction [F(48,400)=3.69; *p*<0.0001]. Post hoc analysis within group confirmed that significantly more infusions were obtained compared to session 1 by LgA rats (sessions 6-17), norBNI+LgA rats (sessions 13, 16 and 17), and LgA-7day rats (sessions 8-17), whereas self-administration did not change for the saline group (N=6). The number of infusions by LgA rats was significantly higher than all other groups during sessions 11, 13, 16 and 17. Post-hoc analysis confirmed significant increases in infusions following the first, second and third weekend breaks for the LgA group (**Fig. 4B**) but there were no significant differences for the other three treatment groups across the same sessions.

**Figure 4.**
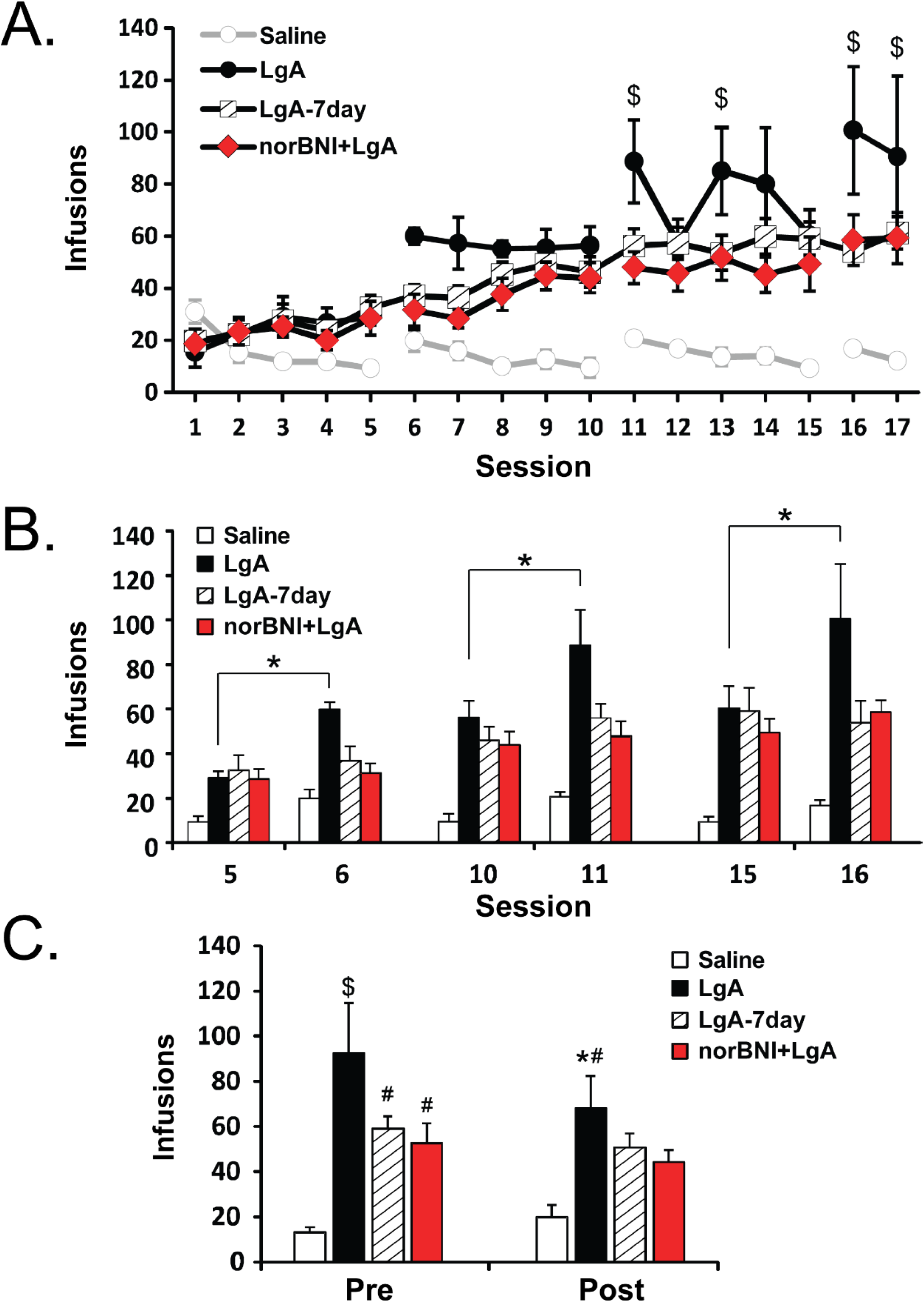
Manipulation of KOR signaling through daily access or systemic administration of norBNI ameliorates escalation of oxycodone self-administration. Mean (±SEM) infusions for groups of male rats trained to self-administer oxycodone (0.15 mg/kg/inf) during weekdays (LgA, norBNI+LgA) or daily sessions (LgA-7day) during A) acquisition, B) pre- and post-weekends, and following 30-day abstinence (reflects averages of the 4 sessions immediately before (Pre) and after (Post) the abstinence period). Broken lines between sessions 5 & 6, 10 &11 and 15 & 16 represent 60h withdrawal period. Significant differences within-group across time are indicated by * and between group compared with saline vehicle condition by # and differences from all other groups by $ for given session.

Following the 30-day abstinence period, there was no significant change in oxycodone infusions obtained under a FR1 response contingency in saline, LgA-7day and norBNI+LgA rats, whereas intake was decreased in LgA rats (**Fig. 4C**). One rat (LgA) that completed the acquisition interval did not complete the post-30 day abstinence due to catheter loss. The rmANOVA confirmed a significant main effect of Drug Condition [F(3, 24)=8.19; *p*<0.001], of Time [F(3, 24)=7.03; *p*<0.05], and of the Drug Condition x Time interaction [F(3, 24) = 3.28; *p*<0.05]. Post-hoc analysis confirmed that significantly more infusions were obtained by LgA, LgA-7day, and norBNI+LgA rats compared to saline controls pre-abstinence, and LgA-7day and norBNI+LgA rats obtained significantly fewer infusions than LgA rats. After the extended 30-day abstinence, only LgA rats had significantly higher intake compared to saline controls. Within-group comparison showed LgA decreased intake post-abstinence, which may be attributable to loss of functional tolerance following cessation.

### Blockade of KOR signaling, or uninterrupted daily access, attenuates neuroadaptations in the CeA associated with long-access oxycodone self-administration

We then assessed whether continuous daily sessions of oxycodone IVSA or systemic norBNI pretreatment would reverse the neuroadaptations at CeA GABAergic synapses induced by long access to oxycodone. Thus, we compared the effects of U-50488 and norBNI on spontaneous GABA transmission in neurons from the LgA-7day and norBNI+LgA rats to those observed in the LgA group (**Fig. 5**). In the LgA-7day group, 1 μM U-50488 caused a significant decrease in mIPSC frequency [t(6)=3.81, *p*<0.01] to 77.32±5.96% of baseline (**Fig. 5C**). In the norBNI+LgA group, U-50488 also significantly decreased the mIPSC frequency [t(13)=3.59, *p*<0.01] to 82.0±5.01% of baseline, but also caused a small but significant increase in amplitude [t(13)=3.52, *p*<0.01] to 109.2±2.6% of baseline and a significant increase in decay time [t(13)=4.05, *p*<0.01] to 125.4±6.27% of baseline. A one-way ANOVA of the effects of U-50488 on each mIPSC characteristic between LgA, LgA-7day, and norBNI+LgA groups confirmed a significant difference in the effects on mIPSC amplitude [F(2,33)=7.53, *p*<0.01] (with a Bonferroni post-hoc confirming differences between the LgA group and both the LgA-7day group [t(33)=2.92, *p*<0.05] and the norBNI+LgA group [t(33)=3.50, *p*<0.01]). Bath application of norBNI had no effect on the baseline mIPSC characteristics of LgA-7day or norBNI+LgA rats (**Fig. 5D**). However, one-way ANOVAs comparing the effects of norBNI for each mIPSC characteristic between these groups and the LgA group confirmed significant differences in the effects of norBNI on mIPSC frequency [F(2,34)=3.65, *p*<0.05], amplitude [F(2,34)=9.47, *p*<0.001], and decay time [F(2,34)=4.44, *p*<0.05]. Bonferroni post-hoc tests further confirmed significant differences between LgA and LgA-7day groups for frequency [t(34)=2.63, *p*<0.05] and amplitude [t(34)=3.99, *p*=0.001], and significant differences between LgA and norBNI+LgA groups for amplitude [t(34)=3.53, *p*<0.01] and decay time [t(34)=2.97, *p*<0.05].

**Figure 5.**
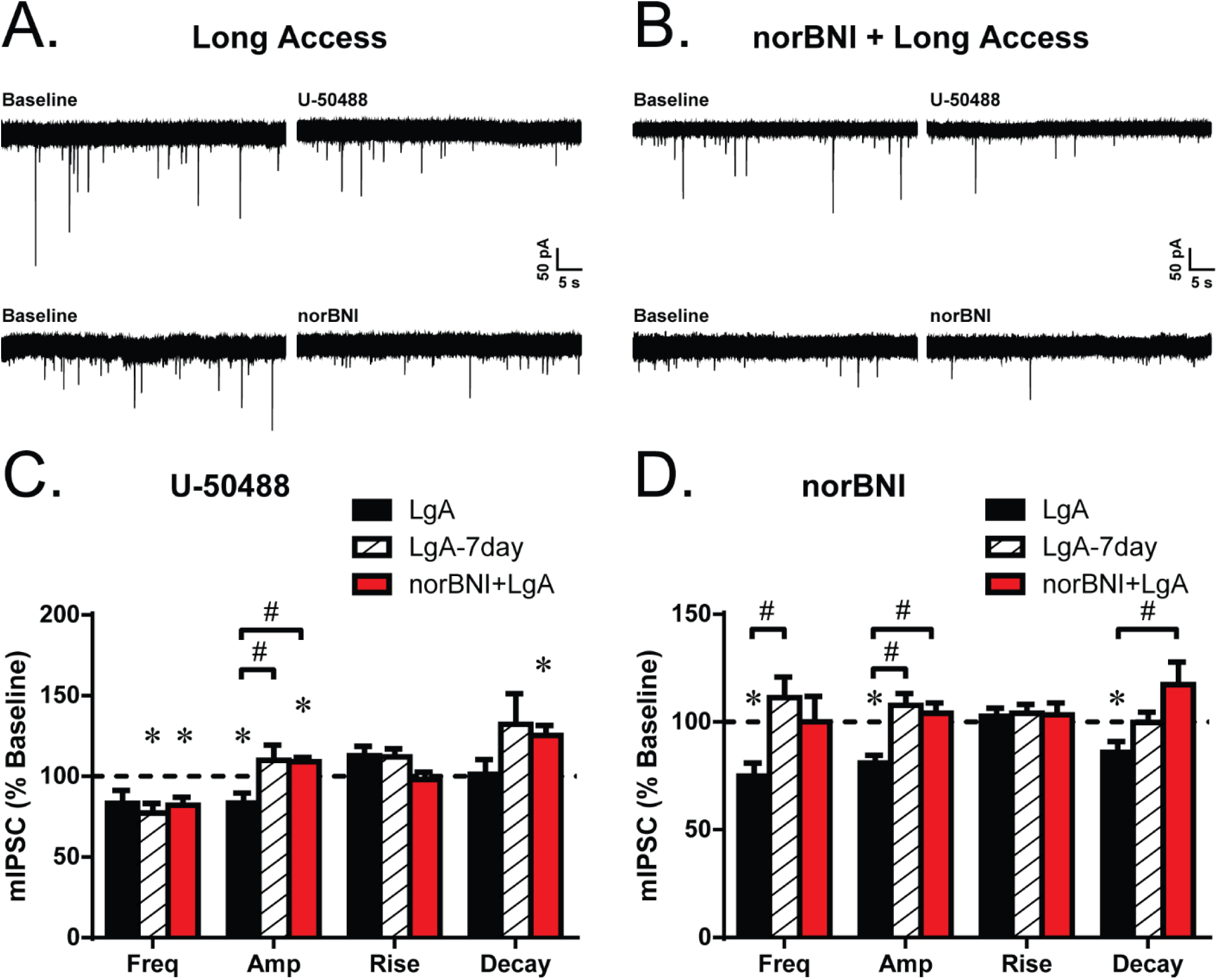
Manipulation of KOR signaling through daily access or systemic administration of norBNI ameliorates oxycodone extended access effects in the CeA. Representative mIPSCs recorded before and after U-50488 (1 μM) or norBNI (200nM) application in neurons from A) LgA and B) norBNI+LgA rats. Mean effect of D) U-50488 or E) norBNI on mIPSC frequency, amplitude, rise time, and decay time in CeA neurons from LgA, LgA-7day, and norBNI+LgA rats. * denotes significant effect of drug compared to baseline and # denotes significant difference between groups.

### Extended access to oxycodone self-administration decreases dynorphin expression in the CeA

Brain tissue were collected from a subset of the LgA-7day rats (N=7; not used for electrophysiology experiments) and sections containing CeA were analyzed for dynorphin A immunoreactivity and compared to sections from naïve control rats (N=8). As shown in **Fig. 6**, oxycodone self-administration under long-access conditions significantly decreased the number of dynorphin A cells in the CeA compared to naïve controls [t(13)=2.384, *p*<0.05], indicating a downregulation of dynorphin peptide that is consistent with lowered endogenous dynorphin activity. This result is also consistent with the lack of effect of KOR antagonism on mIPSCs in this group. This confirmed that escalated IVSA of oxycodone dysregulates the endogenous KOR ligand.

**Figure 6.**
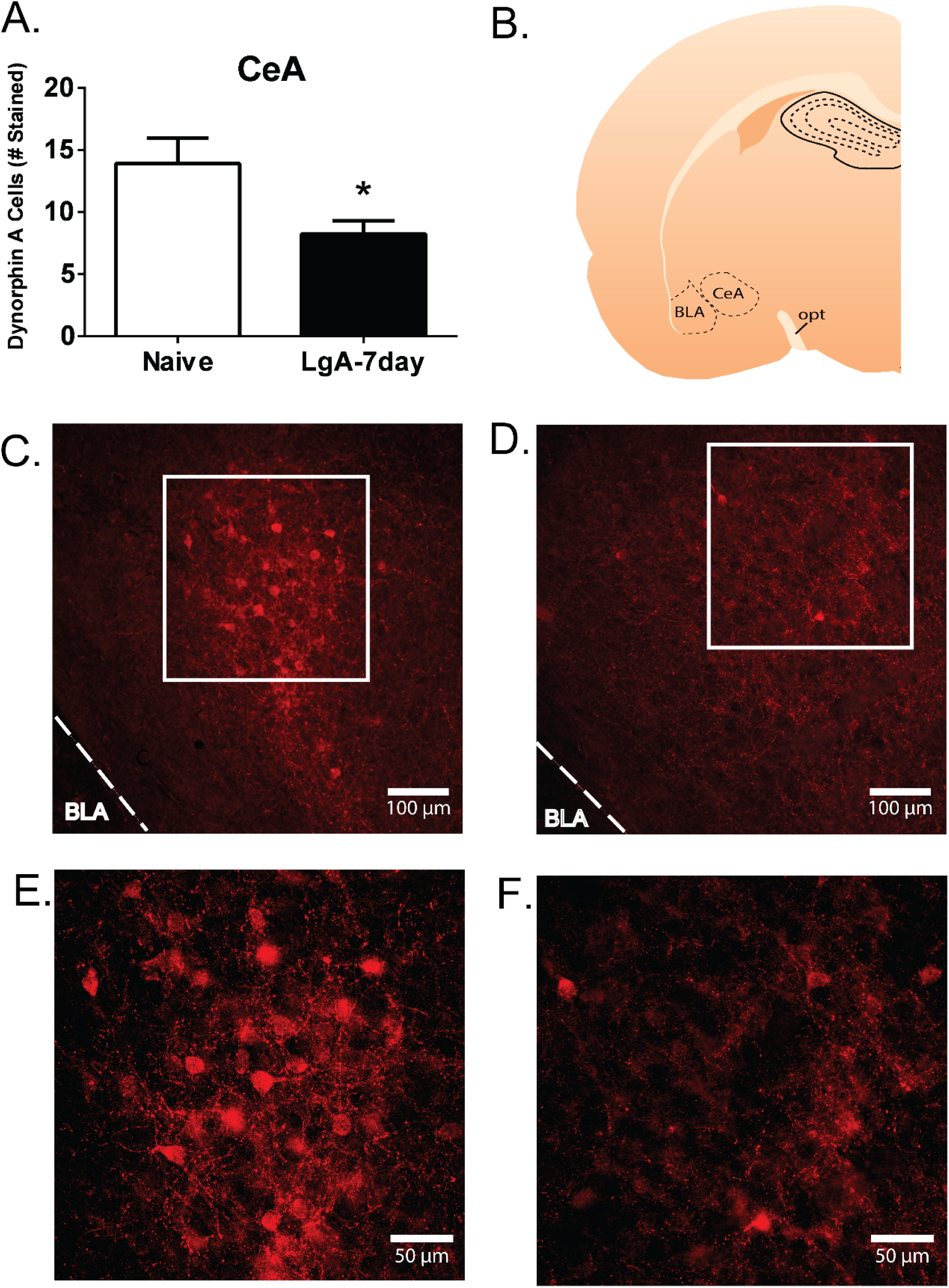
Extended access to oxycodone self-administration decreases dynorphin expression in the CeA. A) Significant reduction in the number of dynorphin A-immunoreactive cell bodies in the CeA of Long access (LgA-7day) rats (N=7) as compared with naïve rats (N=8). B). Example of the approximate coronal section corresponding with the images in panels C-F. Representative 10X photomicrographs show a qualitative decrease in observable dynorphin A cell bodies in LgA-7day rats D) as compared with naïve rats C). E) 20X photomicrograph showing magnified view of the field with in the box in panel C) and F) depicts the magnified field from panel D). Scale bars = 100 μm in C&D), 50 μm in E&F). Data are shown as mean ± SEM. \**p*=0.033. BLA, basolateral amygdala; opt, optic tract.

## DISCUSSION

Nonmedical use of prescription opioid medications has increased tremendously in the US over the past two decades. Oxycodone is one of the most common drugs diverted for nonmedical use, yet there has been relatively little preclinical investigation into the consequences of such use. This dearth of information is particularly acute with respect to both laboratory animal IVSA procedures, the gold standard for assessing abuse liability, and the determination of cellular mechanisms and synaptic effects of oxycodone abuse. One prior investigation determined that long (12h) daily access to oxycodone in IVSA procedures resulted in progressively escalated drug taking compared with 1h access (8). In this study, we confirm that animals provided access to oxycodone in 12h sessions increase their mean intake, whereas animals allowed 1h access sessions do not. We also determined that intermittently increasing the interval of withdrawal directly leads to further increased oxycodone self-administration which is at least in part due to disruption of KOR-mediated regulation of GABA function in the CeA.

The negative reinforcement hypothesis suggests that daily withdrawal from drug access grows increasingly dysphoric and aversive, thus explaining the gradual escalation of drug self-administration. However, this hypothesis has typically only been inferred from studies in which only fixed withdrawal intervals from the end of the previous session to the beginning of the next day’s session are included. The role of varying intervals of withdrawal in the course of escalation has not been explicitly tested, although it is noted that several prior studies report 5-7 day per week scheduling of opioid IVSA sessions (8-10). The explicit comparison of Monday-Friday (M-F) versus 7 days per week scheduling of IVSA sessions in the present study confirms that extended 60 h deprivations increase oxycodone IVSA, resulting in a step-wise, upward ratcheting pattern of behavior, in contrast to the gradually-increasing pattern of intake observed in the 7 days per week group (12h deprivations). Furthermore, this upward-ratcheting pattern depended on intact KOR function since systemic pretreatment with the long-lasting KOR antagonist norBNI resulted in a pattern of IVSA in animals scheduled M-F that was identical to the ones scheduled 7 days per week.

Additionally, dynorphin immunoreactivity in the CeA of animals from the LgA-7day group was significantly lower than that of naïve controls. As the escalation of the LgA-7day group was very similar to that of the norBNI+LgA group, it is likely that dynorphin/KOR signaling is impaired in a similar fashion between these groups. This suggests that long access oxycodone self-administration is sufficient to decrease endogenous dynorphin signaling in the CeA. This result is somewhat surprising given that extended access to heroin has been shown to increase prodynorphin immunoreactivity in nucleus accumbens (9). Presumably, increasing withdrawal severity would increase dynorphin/KOR signaling, driving increased intake of oxycodone to relieve dysphoria. This may underlie the upward-ratcheting pattern of intake following the 60h withdrawal seen in the LgA group. Thus, the time course of dynorphin expression may initially decrease following oxycodone self-administration, and ramp up after sufficient withdrawal time (60h).

This is the first study to examine synaptic effects of withdrawal-induced neuroadaptations in the CeA during extended access oxycodone self-administration. Interestingly, we found no baseline differences between saline, ShA, and LgA animals in both evoked and spontaneous action potential-independent CeA GABAergic signaling, which contrasts with effects found following extended access to IVSA of other drugs of abuse. Extended cocaine access did not affect basal spontaneous action potential-independent CeA GABAergic transmission, but it did increase basal evoked and spontaneous action potential-dependent GABAergic transmission, suggesting increased GABA release (23, 27). This increase is also seen in the CeA of alcohol dependent rats (38), but not in chronic morphine treated rats (39). Thus, despite chronic use of other drugs causing neuroadaptations to opioid receptor systems in the CeA (23, 40), activation of opioid receptors does not appear to alter their role in basal GABAergic neurotransmission.

In contrast, extended access oxycodone self-administration altered the sensitivity of the CeA GABAergic synapses to exogenous KOR ligands. Interestingly, KOR-mediated effects on evoked GABAergic signaling were unaffected by oxycodone self-administration, but spontaneous action potential-independent GABA release was altered, suggesting synapse specific effects. While the KOR agonist U-50488 decreased GABAergic transmission similarly in the saline, ShA and LgA groups, the effects of acute application of the KOR antagonist norBNI varied between these groups. NorBNI caused an increase in GABA release in saline animals, indicating the presence of tonic dynorphin/KOR signaling, but this effect was abolished in ShA animals in an intermediate phenotype and reversed in LgA animals. This “switch” in the effect of norBNI was also reported for extended access cocaine animals (23), although that was accompanied by a switch in the effects of U-50488 on GABAergic signaling. KORs, being G-protein coupled receptors, can activate a multitude of downstream signaling cascades that generally lead to an inhibition of neurotransmitter release, but many of these signaling pathways may be KOR ligand specific (41). In fact, it has been proposed that the long-lasting KOR inhibitory effects of norBNI are due to activation of the c-Jun N-terminal Kinase (JNK) pathway, in a manner distinct from JNK activation through U-50488- or dynorphin-activated KORs (42). Thus, extended access oxycodone administration may selectively dysregulate this ligand-specific KOR-JNK pathway resulting in the specific switch seen with acute norBNI application, while leaving dynorphin/KOR agonist signaling intact. This norBNI-JNK pathway is not well understood (41, 43), thus further investigation into the involvement of dysregulated JNK signaling is warranted.

The synaptic effects were mirrored in the behavioral results, where neither the 7-day per week group nor the norBNI pretreatment group exhibited the norBNI switch seen with the LgA group. Interestingly, the effects of U-50488 and norBNI in the LgA-7day and norBNI+LgA groups looked remarkably similar to the effects of these drugs in the ShA group, despite the drastic increase in oxycodone intake throughout the experiment. The lack of effect of norBNI on these cells would suggest that the ShA group may also have decreased dynorphin expression as seen in the LgA-7day group. Interestingly, *Pdyn* mRNA expression was decreased in the periamygdaloid-cortex of rats 24h after a short access paradigm (3h) of heroin self-administration, and this decrease was consistent in postmortem brain tissue of human heroin addicts (44). For the norBNI+LgA group, the long-lasting activation of the KOR-JNK pathway to inhibit other KOR signaling may have prevented oxycodone-induced dysregulation of this pathway. In the LgA-7day group, this would suggest that increasing withdrawal severity was necessary to induce the proposed dysregulation in the KOR-JNK pathway. Interestingly, withdrawal from chronic alcohol use decreased JNK mRNA expression in the CeA (45).

So called “deprivation effects” have been previously reported for the self-administration of drugs of abuse. This generally refers to an increase in alcohol or drug consumption after an abstinence interval greater than the usual day-to-day interval. For example, Heyser and colleagues showed that ethanol intake increases after a period of 3-28 days of deprivation compared to baseline (46). Such studies have rarely manipulated the deprivation interval in the course of initial escalation training to determine if the rate of escalation is affected by increasing one or more of the inter-session interval(s). Intermittent periods (24-48h) of abstinence produced escalation of nicotine intake under extended access conditions (47). An increase in nicotine infusions compared to baseline was observed following a single deprivation period, however a pattern of further increase during subsequent sessions preceded by 24-48h deprivation periods was not observed. Similarly, intermittent access followed by 7 day abstinence during the early stages of cocaine abuse significantly potentiated drug seeking (48). It is therefore not possible to conclude if this reflects a unique property of oxycodone within the class of opioids or if similar patterns might be observed with other drugs of abuse (e.g., psychostimulants, heroin) with the appropriate experimental design.

Importantly, these findings may have clinically relevant implications. The highest escalation of oxycodone self-administration, as well as the greatest dysregulation of KOR function in the CeA, was found in the group that had extended access to oxycodone 5 days per week with intermittent longer withdrawal periods. Therefore, these data would suggest that individuals on short-term prescribed oxycodone regimens may not wish to skip prescribed days of treatment in an attempt to ‘tough it out’, as this may in fact lead to increased liability for abuse. These findings highlight the importance of adherence monitoring or adherence enhancing interventions, as non-adherence to pain medication use is very common (49) and particularly because there does not appear to be an association between medication adherence and pain treatment outcome (50).

Overall, we conclude that access duration differentially impacts the acquisition and maintenance of the self-administration of oxycodone and that escalation of drug intake is mediated by the negative affective state associated with oxycodone withdrawal, as demonstrated by KOR-induced dysregulation of GABAergic signaling in the CeA. These data further suggest that timing of medication administration may be a critical factor during the early stages of oxycodone addiction.

## ACKNOWLEDGMENTS

This work was funded by support from the United States Public Health Service National Institutes of Health grants R01s DA035281 to M.A.T. and AA015566 to M.R., P60 AA06420 to M.R., F32 AA025262 to D.K., and T32 AA007456 to D.K., M.Q.S., and R.P., as well as the Austrian Science Fund (FWF J 3942-B30 to S.K.), and the Pearson Center for Alcoholism and Addiction Research, all of which had no direct input on the design, conduct, analysis or publication of the findings. We thank Sarah A. Laredo for help with figure diagrams. This is manuscript #29556 from The Scripps Research Institute.

## SUPPLEMENTAL MATERIALS FOR

Withdrawal-induced escalation of oxycodone self-administration is mediated by kappa opioid receptor function

### SUPPLEMENTAL METHODS

#### Intravenous catheterization

The intravenous catheters consisted of a 14.5-cm length of polyurethane based tubing (Micro-Renathane^®^, Braintree Scientific, Inc, Braintree, MA) fitted to a guide cannula (Plastics One, Roanoke, VA) curved at an angle and encased in dental cement anchored to an ∼3 cm circle of durable mesh. Catheter tubing was passed subcutaneously from the animal’s back to the right jugular vein. Catheter tubing was inserted into the vein and tied gently with suture thread. A liquid tissue adhesive was used to close the incisions (3M™ Vetbond™ Tissue Adhesive: 1469SB, 3M, St. Paul, MN). A minimum of 4 days was allowed for surgical recovery prior to starting an experiment. For the first three days of the recovery period, an antibiotic (cefazolin) and an analgesic (flunixin) were administered daily. During testing and training, intravenous catheters were flushed with ∼0.2-0.3 ml heparinized (166.7 USP/ml) saline before sessions and ∼0.2-0.3 ml heparinized saline containing cefazolin (100 mg/mL) after sessions.

Catheter patency was assessed via administration through the catheter of ∼0.2 ml (10 mg/ml) of the ultra-short-acting barbiturate anesthetic Brevital sodium (1% methohexital sodium; Eli Lilly, Indianapolis, IN). Animals with patent catheters exhibit prominent signs of anesthesia (pronounced loss of muscle tone) within 3 sec after infusion. Animals that failed to display these signs were considered to have faulty catheters and were discontinued from the study.

#### Self-administration procedure

##### Acquisition

Drug self-administration was conducted in operant boxes (Med Associates, Inc., Fairfax, VT) located inside sound-attenuating chambers located in an experimental room (ambient temperature 22±1 °C; illuminated by red light) outside of the housing vivarium. To begin a session, the catheter fittings on the animals’ backs were connected to polyethylene tubing contained inside a protective spring suspended into the operant chamber from a liquid swivel attached to a balance arm. Each operant session started with the extension of two retractable levers into the chamber. Following each completion of the response requirement (response ratio), a white stimulus light (located above the reinforced lever) signaled delivery of the reinforcer and remained on during a 20-sec post-infusion timeout, during which responses were recorded but had no scheduled consequences. Drug infusions were delivered via syringe pump. The training dose (150 μg/kg/infusion; ∼0.1 ml/infusion) was selected from prior self-administration studies (1). In Behavior Experiment 1, session duration for the normal (Short Access; ShA, N=12) acquisition was 1h and the Long Access (LgA, N=12) training sessions were 12h in duration. Rats received access for 4 sequential sessions in week 1, 5 sessions/week thereafter (i.e., with 60h weekend abstinence periods). Two rats in the LgA group that failed to average more than 24 infusions (defined by the highest responder in the Saline group, see below) over the final 5 sessions of acquisition were excluded from the acquisition and PR self-administration data as non-escalators. Two additional rats in the LgA group failed patency testing during the acquisition interval and were excluded. One additional rat from the LgA group was euthanized during abstinence due to illness. In Behavior Experiment 2, the session duration was 12h for 5 sessions/week for a LgA oxycodone group (N=6) and a group responding only for i.v. Saline (N=6). One rat from the LgA group had its catheter rendered nonfunctional by the cage mate after 15 sessions, therefore the session 15 value was used for the final two sessions for this animal. A third group was permitted to respond for oxycodone for consecutive daily 12h sessions (LgA-7day; N=12); one rat failed patency during the abstinence interval and was excluded. A fourth group was treated with norBNI (30 mg/kg, i.p.) and then trained for 5 LgA sessions/week (norBNI+LgA; N=6). Rats were returned to their home cages for an extended 30-day abstinence period and then returned to 12h sessions to test for re-engagement of drug seeking (re-escalation) following detoxification.

##### Progressive-Ratio (PR) Dose-Response Testing

For the PR task, the sequence of response ratios started with one response then progressed thru ratios determined by the following equation (rounded to the nearest integer): response ratio=5e^(injection number x j) – 5 (2). The value of ‘j’ was 0.2 and was chosen so as to observe a ‘breakpoint’ within ∼3h. The last ratio completed before the end of the session (1h after the last response up to a maximum 30h sessions) was operationally defined as the breakpoint. The rats completed series in which oxycodone (0, 60, 150, 300 μg/kg/inf) was available, and the dose order was balanced by Latin Square design (i.e. the 4 total conditions were randomized). Session duration was up to 3h.

#### Electrophysiology

##### Slice Preparation

Rats were anesthetized with isoflurane (3-5%) followed by rapid decapitation and removal of the brain immediately into an ice-cold high sucrose cutting solution (sucrose 206 mM; KCl 2.5 mM; CaCl_2_ 0.5 mM; MgCl_2_ 7mM; NaH_2_PO_4_ 1.2 mM; NaHCO_3_ 26 mM; glucose 5 mM; HEPES 5 mM; pH 7.4). CeA coronal slices (300-400 μm) were cut on a Leica 1200S vibratome (Buffalo Grove, IL), incubated in an interface configuration for 30 min to 1 h, and then submerged and continuously superfused (flow rate of 2-4 ml/min) with 95% O_2_/5% CO_2_ equilibrated artificial cerebrospinal fluid (aCSF) of the following composition: NaCl 130 mM; KCl 3.5 mM; NaH_2_PO_4_ 1.25 mM; MgSO_4_·7H_2_O 1.5 mM; CaCl_2_ 2.0 mM; NaHCO, 24 mM; glucose 10 mM. The number of cells in each experimental group were from a minimum of four animals.

##### Intracellular Recordings of Evoked IPSPs

We recorded from neurons in the medial subdivision of the CeA with sharp micropipettes filed with 3M KCl using discontinuous current-clamp mode. Neurons were held near their resting membrane potential (RMP). Data were acquired with an Axoclamp 2B amplifier (Axon Instruments, Foster City, CA) and stored for later analysis using pClamp software (Molecular Devices, Sunnyvale, CA). We evoked GABAergic inhibitory postsynaptic potentials (IPSPs) by stimulating locally within the CeA through a bipolar stimulating electrode. We performed an input–output (I/O) protocol consisting of a range of five current stimulations, starting at the threshold current required to elicit an IPSP, up to the strength required to elicit the maximum amplitude. These stimulus strengths were maintained throughout the duration of the experiment. We examined paired pulse ratio (PPR), where two stimuli of equal intensity were applied at an interstimulus interval of 50 ms (3). The PPR is calculated as the ratio of the amplitude of the second IPSP over the first IPSP, with stimulus intensity set such that the first IPSP amplitude was ∼50% of the maximum amplitude determined by the I/O protocol. A drug induced change in PPR suggests presynaptic effects of the drug, where an increase in PPR reflects a decrease in neurotransmitter release. All measures were performed prior to (baseline) and during drug application.

##### Whole-Cell Patch Clamp Recording of miniature IPSCs

We recorded from medial CeA neurons visualized in brain slices using infrared differential interference contrast (IR-DIC) optics and CCD cameras (QImaging, Surrey, BC, Canada). Whole-cell voltage-clamp recordings were made with a Multiclamp 700B amplifier (Molecular Devices), low-pass filtered at 10 kHz, digitized (Digidata 1440A and 1550B; Molecular Devices), and stored on a PC using pClamp 10 software (Axon Instruments). All voltage clamp recordings were performed in a gap-free acquisition mode, with cells clamped at –60 mV for the duration of the recordings. Patch pipettes (3-6 MΩ) were pulled from borosilicate glass (Warner Instruments, Hamden, CT; King Precision, Claremont, CA) and filled with an internal solution composed of (in mM): 145 KCl; 0.5 EGTA; 2 MgCl_2_; 10 HEPES; 2 Na-ATP; 0.2 Na-GTP. GABAergic miniature IPSCs (mIPSCs) were recorded in the presence of 1 μM tetrodotoxin (TTX). In all experiments, series resistance was continuously monitored with a 10 mV pulse and neurons with >20% change in series resistance were not included in the final analysis. All measures were performed prior to (baseline) and during drug application.

#### Immunohistochemistry

LgA-7day rats (N=7) and naïve controls (N=8) were euthanized by decapitation and brains were rapidly removed and fixed overnight in 5% acrolein in PBS overnight at 4°C. The following day, brains were cryoprotected in 25% sucrose in PBS at 4°C until they sank, were flash frozen in pre-chilled isopentane on dry ice, and stored at - 80°C until cryosectioning. All sections were immunostained in a single batch. Brains were sectioned at 40 μm and free floating sections were washed twice for 5 min in PBS. Sections were subsequently incubated for 10 min in 0.1 M sodium borohydride in PBS, treated for 30 min in a 1:1 mixture of ethanol and PBS containing 0.3% hydrogen peroxide, and blocked for 2 hr in 10% normal donkey serum (NDS) in PBS. Sections were incubated 48 h at 4 °C in rabbit anti-dynorphin antibody (1:1500, H-021-03, Phoenix Pharmaceuticals, Burlingame, CA) diluted in PBS with 0.5% Triton X (PBS-Tx) with 2% NDS. Following primary incubation, tissue was washed three times for 5 min each in PBS (3× 5 min-washes) and then incubated for 1.5 h at room temperature in biotinylated-donkey anti-rabbit IgG (1:750, 711-065-152, Jackson Immunoresearch, West Grove, PA) in PBS-Tx with 2% NDS. Sections were washed 3x5 min in PBS and incubated 30 min at room temperature in an Alexa Fluor-647 conjugated streptavidin (1:500, S21374, Invitrogen, Waltham, MA). Tissue was then washed overnight at 4 °C in PBS. Stained sections were mounted onto Superfrost plus slides (Thermo Fisher Scientific, Waltham, MA) and coverslipped using ProLong Gold antifade medium (Thermo Fisher Scientific). Antibody specificity was tested by omission of primary which yielded no staining. This antibody has previously been used in rat (4, 5), and presadsoprtion with dynorphin A1-17 blocked all staining while preadsorption with β-endorphin and leu-enkephalin had no effect (5).

Sections containing central amygdala were stained every 120 μm beginning from approximately Bregma level -1.92 for a total of four sections. A monochromatic camera connected to a Keyence BZX digital scope was used to image slides. Using ImageJ software (NIH, Bethesda, MD) a 0.7 x 0.55 mm box was placed in the lateral central amygdala. The number of dynorphin-immunoreactive perikarya were manually counted by an observer who was blinded to treatment. Bilateral measurements were averaged together for each section and then sections were averaged together to obtain average cell count per brain.

#### Data Analysis

Analyses of behavioral data and immunohistochemistry were conducted using Prism for Windows (v.5, v.6.02; Graphpad Software, Inc, La Jolla, CA). Graphs were generated with Prism and Excel (Microsoft, Redmond, WA) and figures created in Canvas (v.12, v.16; ACD Systems of American, Inc, Seattle, WA) and Adobe Illustrator CS6 (v.16.0.3, Adobe Systems Inc, San Jose, CA).

**SUPPLEMENTAL FIGURE 1.**
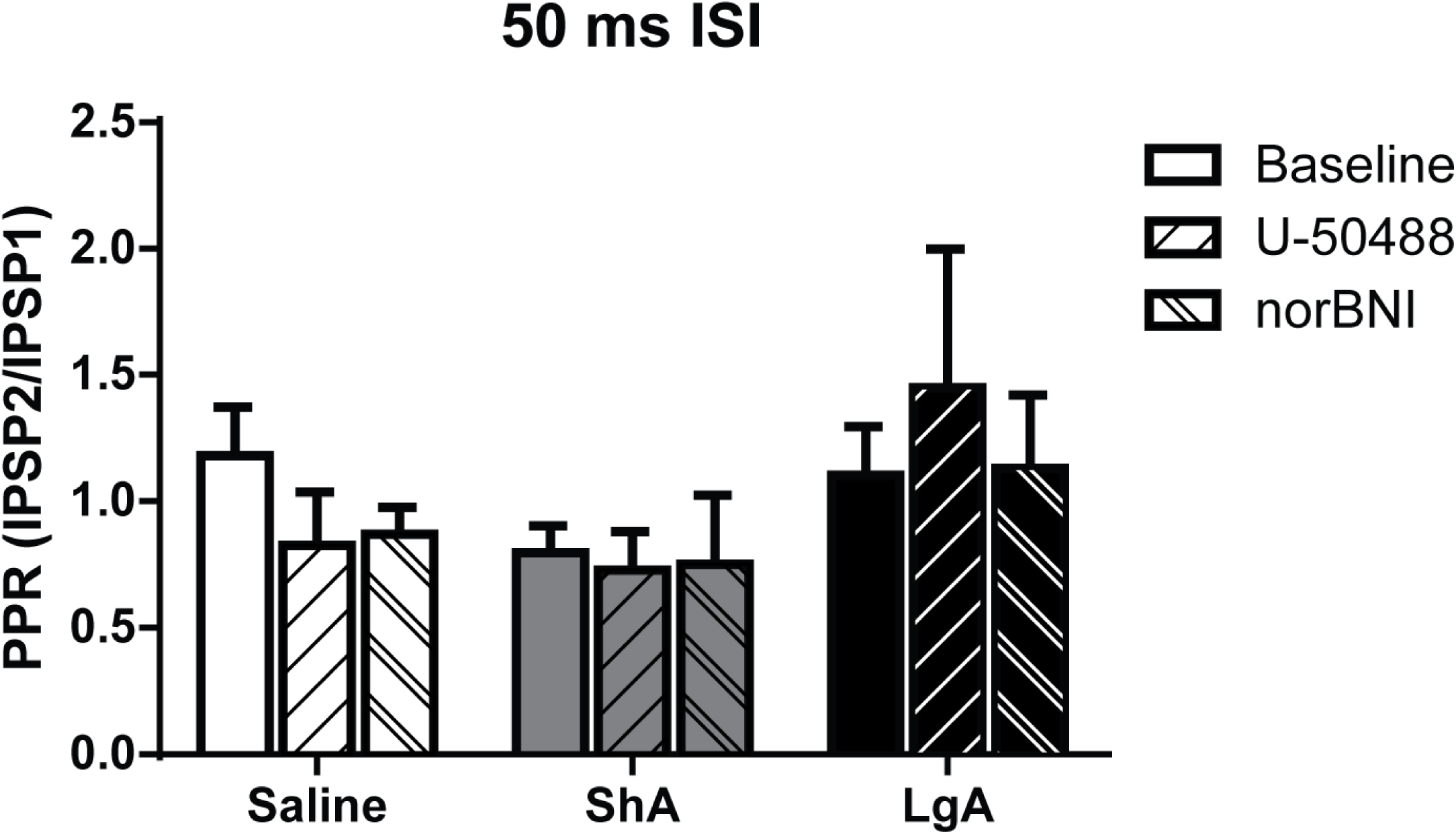
Paired pulse ratio (PPR) of CeA neurons in slices from Saline, ShA, and LgA animals at an insterstimulus interval (ISI) of 50 ms. There was no significant difference between baseline PPR among Saline, ShA, and LgA animals, and no significant effect of either U-50488 or norBNI on baseline PPR among Saline, ShA, or LgA animals.

